# BiG-MAP: an automated pipeline to profile metabolic gene cluster abundance and expression in microbiomes

**DOI:** 10.1101/2020.12.14.422671

**Authors:** Victoria Pascal Andreu, Hannah E. Augustijn, Koen van den Berg, Justin J. J. van der Hooft, Michael A. Fischbach, Marnix H. Medema

## Abstract

Microbial gene clusters encoding the biosynthesis of primary and secondary metabolites play key roles in shaping microbial ecosystems and driving microbiome-associated phenotypes. Although effective approaches exist to evaluate the metabolic potential of such bacteria through identification of metabolic gene clusters in their genomes, no automated pipelines exist to profile the abundance and expression levels of such gene clusters in microbiome samples to generate hypotheses about their functional roles and to find associations with phenotypes of interest. Here, we describe BiG-MAP, a bioinformatic tool to profile abundance and expression levels of gene clusters across metagenomic and metatranscriptomic data and evaluate their differential abundance and expression between different conditions. To illustrate its usefulness, we analyzed 47 metagenomic samples from healthy and caries-associated human oral microbiome samples and identified 58 gene clusters, including unreported ones, that were significantly more abundant in either phenotype. Among them, we found the *muc* operon, a gene cluster known to be associated to tooth decay. Additionally, we found a putative reuterin biosynthetic gene cluster from a *Streptococcus* strain to be enriched but not exclusively found in healthy samples; metabolomic data from the same samples showed masses with fragmentation patterns consistent with (poly)acrolein, which is known to spontaneously form from the products of the reuterin pathway and has been previously shown to inhibit pathogenic *Streptococcus mutans* strains. Thus, we show how BiG-MAP can be used to generate new hypotheses on potential drivers of microbiome-associated phenotypes and prioritize the experimental characterization of relevant gene clusters that may mediate them.

**Importance:** Microbes play an increasingly recognized role in determining host-associated phenotypes by producing small molecules that interact with other microorganisms or host cells. The production of these molecules is often encoded in syntenic genomic regions, also known as gene clusters. With the increasing numbers of (multi-)omics datasets that can help understanding complex ecosystems at a much deeper level, there is a need to create tools that can automate the process of analyzing these gene clusters across omics datasets. The current study presents a new software tool called BiG-MAP, which allows assessing gene cluster abundance and expression in microbiome samples using metagenomic and metatranscriptomic data. In this manuscript, we describe the tool and its functionalities, and how it has been validated using a mock community. Finally, using an oral microbiome dataset, we show how it can be used to generate hypotheses regarding the functional roles of gene clusters in mediating host phenotypes.

## Introduction

Bacteria can produce diverse sets of small molecules that interact with other microbes or with their host. These metabolites include members of both primary and secondary metabolism and cover a wide chemical diversity^1,2^. These pathways and metabolites are often specific to certain strains or species and help them to compete for space and resources^3^, e.g. through antimicrobial, nutrient-scavenging or immunomodulatory activities^4^. The genes that encode these pathways are often physically clustered and are also known as Biosynthetic Gene Clusters (BGCs) or Metabolic Gene Clusters (MGCs)^5,6^—the latter being a broader definition that also includes catabolic pathways. Several studies have indicated metabolites produced from such gene clusters to be the major drivers of specific phenotypic traits; for instance, pseudomonads in the rhizosphere of sugar beet plants were shown to produce the antifungal non-ribosomal peptide (NRP) thanamycin, which protects plants from fungal infections^7^. Another example from primary metabolism is trimethylamine, a diet derived-molecule that is processed by bacteria harboring a gene cluster that includes both *CutC* and *CutD*, and has been associated with an increased risk of suffering from cardiovascular disease^8^. Therefore, mining genomes for BGCs or MGCs enables moving the field towards a deeper understanding of function at the molecular level and determine the role a given microbe plays in the ecosystem^9^.

Several tools have been developed to mine genomes for these gene clusters, like antiSMASH^10^, gutSMASH (https://github.com/victoriapascal/gutsmash/tree/gutsmash/) or DeepBGC^11^. In contrast to other tools for functional profiling of microbial communities, such as HUMAnN2^12^, MetaPath^13^, FMAP^14^ and Metatrans^15^, these do not depend on pathways that are present in reference databases like KEGG^16^ or MetaCyc^17^, which only include pathways for which most or all enzymatic steps have been elucidated. In fact, the majority of gene clusters identified by antiSMASH and many gene clusters predicted by gutSMASH encode pathways for which the catalytic steps, intermediates, and final products are yet unknown. However, known pathways that are encoded by gene clusters can also be reliably detected. The detection of complete gene clusters instead of individual enzyme-coding genes likely decreases false positive detections of enzymes that show sequence similarity to reference enzyme sequences but are part of different functional contexts. For these reasons, identification of gene clusters of known and unknown function provides a useful basis to look for functional explanations of microbiome-associated phenotypes of interest. As phenotypes are often triggered by metabolites at physiologically relevant concentrations, while samples without the phenotype lack these metabolites or have them at lower concentrations, assessing gene cluster abundance and expression levels across samples is crucial to predict associations with the phenotype in question. Another significant advantage of profiling the community by combining different omics data is to prioritize the characterization of putative gene clusters that are highly abundant or expressed in samples of interest and thus, help elucidating novel compounds and their biosynthetic pathways.

Here, we present designed BiG-MAP (Biosynthetic Gene cluster Meta’omics Abundance Profiler), which provides a streamlined and automated process to determine BGC/MGC abundance and expression in bacterial communities by mapping metagenomic and metatranscriptomic reads to gene cluster sequences from reference genomes or metagenomic assemblies. BiG-MAP uses MinHash-based redundancy filtering and groups BGCs into families with BiG-SCAPE^18^ to avoid ambiguous mapping, and uses these to output and visualize profiles of MGC abundance or expression levels across samples. Additionally, it calculates differential abundance or expression using either parametric or nonparametric tests. We validate the tool using simulated metagenomic data and show how MGC abundance and expression levels are accurately recapitulated. Finally, to showcase its usefulness, we applied BiG-MAP on a large publicly available metagenome dataset from the human oral microbiome and describe how it successfully identified gene clusters related to bacteria’s specialized primary and secondary metabolism that are (potentially) relevant for caries development. Among others, this collection includes the previously reported *pdu* and cobalamin gene cluster involved in the reuterin synthesis and the *muc* operon, gene clusters that were predicted by gutSMASH and antiSMASH, respectively. Thus, BiG-MAP suggests new lines to explore further the onset and development of oral cavities.

## Results and discussion

### An approach to map metagenomics and metatranscriptomic reads to gene clusters

BiG-MAP maps shotgun sequencing reads onto gene clusters that have been either predicted by antiSMASH^19^ or gutSMASH (manuscript in preparation). It is a Python-based pipeline, which allows downloading datasets from SRA respository, aligning metagenomic or metatranscriptomic reads to gene clusters detected in reference genome collections or in a metagenomic assembly, providing normalized counts across samples, performing differential analyses, and visualizing the results. The tool requires three main inputs: (1) a gene cluster collection obtained from running any “SMASH-based” algorithm, (2) the meta‘omic dataset in FASTQ or FASTA format or, alternatively, the Sequence Read Archive (SRA) accession numbers to download it, and (3) a metadata file with sample information to segregate them into groups and compare their gene cluster content.

BiG-MAP is composed of four different modules (see Fig. 1): (1) BiG-MAP.family, which performs redundancy filtering on the input collection of predicted gene clusters and provides a set of representative gene clusters for the mapping process. (2) BiG-MAP.download, which uses a list of SRA accession ids to download the shotgun data if present in the SRA database (this step is optional). (3) BiG-MAP.map, which maps reads from the metagenomic or metatranscriptomic samples onto the set of representative gene clusters obtained from BiG-MAP.family. (4) BiG-MAP.analyse, which normalizes the counts for sparsity and sequencing depth, performs differential abundance/expression analysis and visualizes the output.

**Figure 1.**
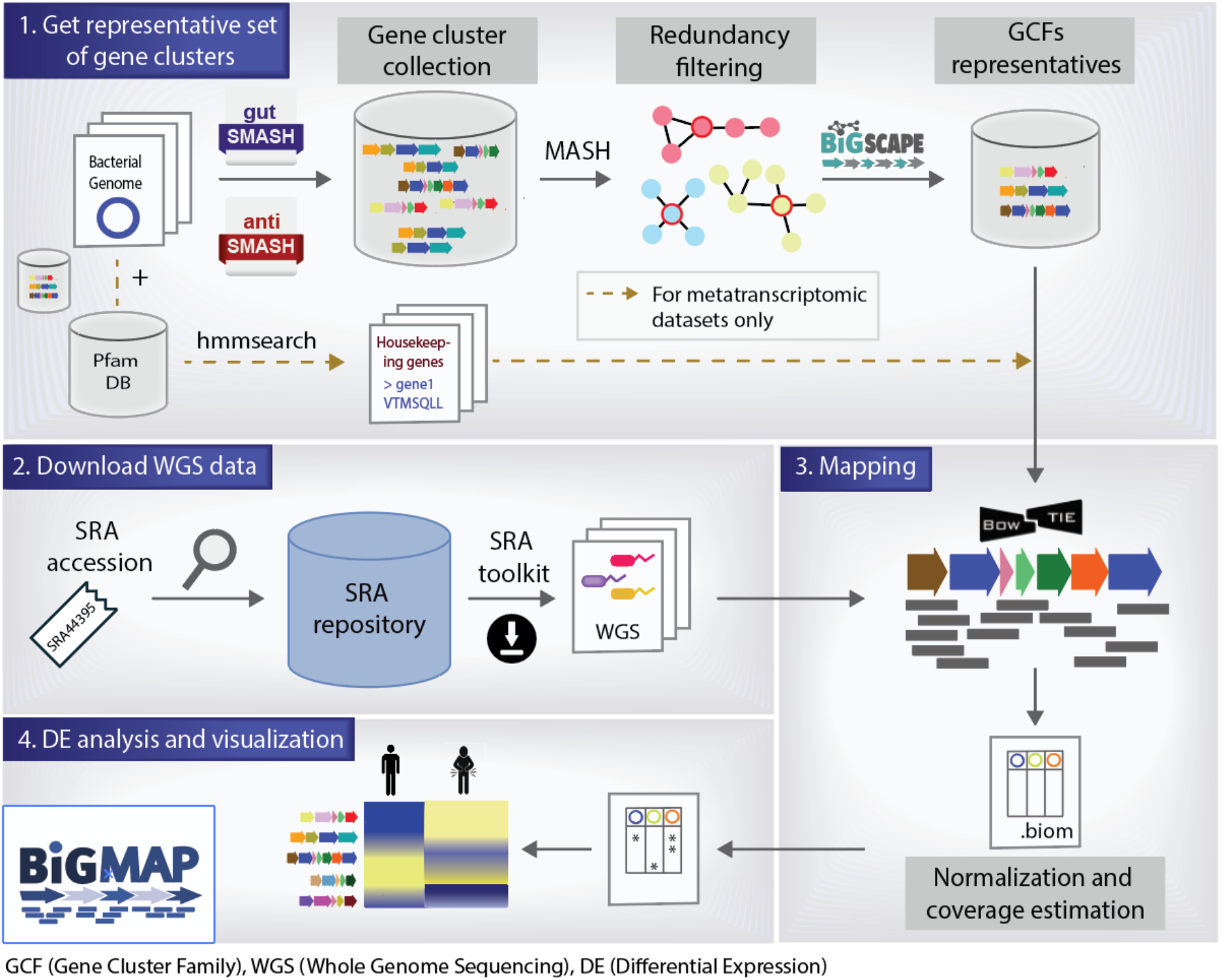
BiG-MAP workflow. BiG-MAP is composed of four different modules: (1) BiG-MAP.family returns a representative set of non-redundant gene clusters based on sequence similarity, given a set of predicted gene clusters by either gutSMASH or antiSMASH. This module also looks for the protein sequences of 5 housekeeping genes from the bacteria encoding the representative gene clusters when reads from metatranscriptomic sequences are going to be used. (2) BiG-MAP.download downloads a set of metagenomes/metatranscriptomes based on their SRA accessions. (3) BiG-MAP.map aligns omics reads to the representative set of gene clusters using Bowtie and (4) BiG-MAP.analyse computes normalized read counts, performs differential abundance/expression analysis of gene clusters across different conditions, and visualizes the results (see Suppl. Fig S1 and S2 as an example).

The BiG-MAP.family module performs a redundancy analysis on the gene cluster collection to remove almost identical sequences, in order to reduce the computing time and avoid ambiguous mapping. To achieve this, the protein sequences of the gene clusters are used as input for MASH^20^, a MinHash-based algorithm to estimate sequence distance. Next, a representative gene cluster is selected using medoids calculation. The resulting representatives are then clustered into Gene Cluster Families (GCFs) using BiG-SCAPE^18^, an algorithm that uses three different distance metrics to group MGCs into families based on sequence and architectural similarity. This step helps to group more distantly related homologous gene clusters that likely have the same chemical products but that are encoded in more distantly related organisms. In such cases, BiG-MAP maps reads to the family representatives separately, but also allows reporting combined abundance or expression levels per family to find associations with phenotypes at a higher level. In order to set an expression baseline when using metatranscriptomic data, BiG-MAP screens bacterial genomes whose gene clusters have been included in the non-redundant representative set of gene clusters for five house-keeping genes known to have stable expression levels using HMMer (for details, see Methods section titled *BiG-MAP.family: Creating a non-redundant MGC representative collection*). Next, the reads are mapped to the representative gene clusters using the short-read aligner Bowtie2^21^. The obtained raw read counts are then converted to RPKM (Reads Per Kilobase Million) values, which are averaged over the GCF size (based on BiG-SCAPE clustering). In the last module, RPKM values are then normalized using Cumulative Sum Scaling^22^ (CSS) to account for sparsity. Moreover, for each aligned gene cluster we assess its coverage to control for gene clusters that are only partially mapped to by meta’omic reads. We report two coverage values in the intermediate files; one for the whole gene cluster and the other considering only the core genes of the BGC/MGC; showing both these numbers is often insightful in cases where borders of gene clusters called by antiSMASH or gutSMASH are imprecise and reads may be mapped to regions flanking the actual gene cluster. Subsequently, BiG-MAP detects differentially abundant or expressed gene clusters by using either zero-inflated gaussian distribution mixture models (ZIG-models) or using a Kruskal-Wallis model. Finally, all the generated results are displayed into a plot that includes a heatmap for the gene clusters abundance/expression values, a bar plot for the log fold change, the coverage values and finally another heatmap for the housekeeping gene expression values when analyzing metatranscriptomes (see Suppl. Fig. S2). The output folders contain different intermediate and final results as for instance the BiG-SCAPE results, the resulting bedgraphs, the raw and normalized RPKM counts for each sample (in BIOM format^23^) and after applying the fitZIG and Kruskal Wallis tests in tab-separated tables and mapping coverage values for each gene cluster and sample. Altogether, this tool presents a streamlined method to functionally profile meta’omics data by mapping reads to known or putative gene clusters.

### Assessing and validating BiG-MAP performance using simulated data

In order to evaluate the overall performance of BiG-MAP and in particular, all the default parameters chosen as defaults, such as the Bowtie alignment mode and the MASH similarity score cut-off, we designed a mock microbial community for metagenome simulation. From the Culturable Genome Reference (CGR) genome collection^24^, we randomly chose 101 CGR genomes to simulate metagenome reads from and to use as input for gutSMASH. To assess the impact of different sequencing depths (coverage of 0.5x and 0.05x) and community structure (uniform, linear, power-law and exponential), we simulated eight different metagenomic libraries. Since the gene cluster content and their abundance levels in simulated data is known (ground truth), this allowed us to assess the recall and precision of the BiG-MAP assignments using MASH dissimilarity scores ranging from 10-100 and the eight different alignment modes available in Bowtie across the eight different simulated data libraries. From these results we computed the F1-score or harmonic mean of precision and recall (see Fig. 2), which showed that the community structure slightly affects BiG-MAP results. Moreover, since the highest F1 scores were obtained when using MASH score cut-off (similarity) of 0.8 and using “fast” alignment mode (end-to-end), we set these parameters as defaults. Still, the user is able to change them as desired by indicating it with the appropriate flag when running BiG-MAP.

**Figure 2.**
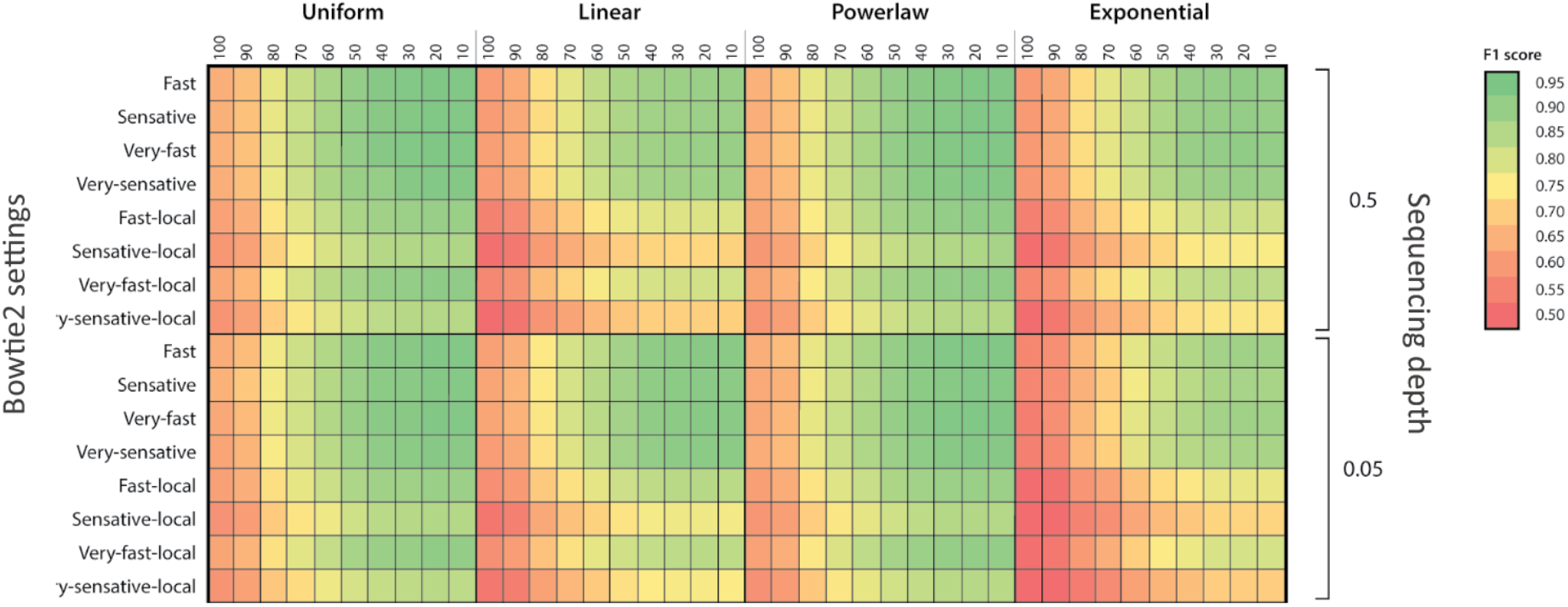
BiG-MAP validation using simulated metagenomes. F1 score heatmap using simulated metagenomes constructed to assess the best MASH dissimilarity cut-off across four different microbial community structures, two different sequencing depth values and eight different Bowtie alignment modes.

### Analysis of the oral microbiome: revealing the presence of gene clusters associated with health and disease

The oral cavity is a natural habitat for many bacteria that reside in or on the gingival sulcus, tongue, teeth and cheeks, among other surfaces. These bacteria take part in important processes such as initial digestion of food, but are also associated with several oral diseases such as caries^25^ and periodontitis^26^. It is known that these bacteria can organize themselves to form biofilms, which can play a causal role in the development of these diseases^27^. There are different functional and metabolic pathway alterations that have been associated with the onset of disease via the production of small molecules^28,29,30,31^.For instance, tetramic acid produced by the caries-associated bacterium *Streptococcus mutans* has been linked to tooth decay^32^. For this reason, in order to functionally profile these oral communities and acquire further insights into the MGCs that might be involved, we studied a dataset of 47 oral microbiome samples^30^ for which paired metagenomics and metabolomics data have been acquired and further analyzed using BiG-MAP (see Methods *Assessing the pdu operon abundance by surveying different oral metagenomic samples* and *Evaluating the presence of the muc operon in caries-associated metagenomes* sections).

To evaluate possible molecular mechanisms underpinning caries formation, we first analyzed the available MS/MS data together with the metabolite feature abundance table using Pathway Activity Level Scoring (PALS)^33^, which uses molecular families obtained using molecular networking^34^ to group similar metabolites, and PLAGE^35^ to find differentially expressed metabolite groups between two conditions. PALS showed a very consistent and strong differential abundance between healthy and caries volunteers of a number of features in a metabolite group that we could annotate with polymer-like structures based on their C_3_H_4_O mass differences. With MASST searches^36^ across all public data present in GNPS-MassIVE, we could confirm the occurrence of these differential features in various microbial, human, and environmental-related public datasets (see Methods and Supplementary Methods for further information on the metabolomics data analysis). Based on the above information, we concluded that these polymer-like structures might well represent molecules called polyacroleins (metabolite identification level 3 - annotated compound class), which are known to spontaneously form from a component of the antimicrobial set of molecules called reuterin^37^, and which have a matching mass difference between different polymer lengths. The formation of (poly)acrolein has been shown to contribute strongly to the antimicrobial activity of reuterin^37^. Reuterin is produced by lactobacilli from a genomic island containing a *pdu-like* operon together with a cobalamin biosynthetic gene cluster^38^. Of note, acrolein is an ubiquitous compound that can be found in the human body for various reasons, such as the endogenous production of it, the ingestion of different food sources or due to exposure to different environmental conditions^39^. There are various known routes that can converge into the formation of acrolein, as it can be formed spontaneously from glycerol and 3-hydroxypropionaldehyde^37^. Furthermore, glycerol metabolism from gut bacteria has also been found to produce this molecule^40^. Typically, the acrolein polymerization occurs under alkaline conditions^41^, thus, it is more likely to accumulate in saliva from healthy samples, as caries typically acidifies the oral cavity. Indeed, our results show that the possible polyacroleins are more abundant in samples of healthy volunteers. Interestingly, the presence of acrolein has been linked to inhibition of *Streptococcus mutans*, a well-known cariogenic bacteria^42,43^.

Based on these findings, we were motivated to look for the presence of the *pdu* operon in the metagenomics samples, in order to identify candidate MGCs that might be involved in acrolein formation. To this end, we ran gutSMASH on the 1,440 genomes from the Human Microbiome Oral Database (HMOD, http://www.homd.org/) available in April 2020. Interestingly, gutSMASH identified a *pdu*-like operon in the genome of *Streptococcus sp. F0442* that also includes a cobalamin (vitamin B12) biosynthetic region and is architecturally similar (cumulative Blast bit score of 13,271) to the *Lactobacillus reuteri* one (see Fig. 3A). Therefore, to assess the abundance of the predicted gene clusters in the oral microbiome we used our gutSMASH run, which predicted 3,352 gene clusters, as input for the BiG-MAP.family module, to filter out redundant MGCs. Next, the reads of the 47 oral metagenomes (24 healthy and 23 caries-related) were mapped onto the 1,544 representative gene clusters using BiG-MAP.map and the counts were further normalized and parsed with BiG-MAP.analyse. We found that 56 gene clusters predicted by gutSMASH were significantly differentially abundant between caries-related and healthy samples when using Kruskal Wallis. Despite the fact that the *pdu* operon was not among these, we could see that it was still somewhat more abundant in healthy samples (mean: 5.30 RPKM counts/sample) when compared to the diseased group (mean: 4.16 RPKM counts/sample). Motivated by this, we sought to assess its presence in a larger oral microbiome dataset by using 48 paired publicly available paired-end metagenome samples, which also included metagenomes from samples suffering from periodontitis and plaque formation, all considered as disease-related samples. These were used in combination with the already analyzed ones, making a total of 96 samples; 33 caries-related, 34 healthy, 10 periodontitis-related and 19 involved in plaque development and all were used as input for BiG-MAP (see Methods section titled *Assessing the pdu operon abundance by surveying different oral metagenomic samples*). From this run, we found 164 gene clusters differentially abundant between groups (using Kruskal Wallis test), and the *pdu* operon was among them. While healthy samples on average have 5.15 RPKM counts/sample mapping to this gene cluster, diseased ones have 3.05 (p-value= 0.0004 using Kruskal Wallis). We also evaluated the coverage of the read mapping within the expanded metagenomic datasets and found that within healthy samples, not all samples contain this gene cluster. For instance, from 34 healthy samples in the extended dataset, we could find 15 of them that appear not to have the *Streptococcus sp. F0442 pdu* operon (coverage below 0.5), while the rest had fairly high coverage scores with a mean coverage value of 0.79 (selecting the samples with coverage values of at least 0.5), implying the presence of this operon or a close homologue of it (see Fig. 3B). Overall, this MGC constitutes a potential source for polyacrolein production, and the hypothesis that it could be involved in inhibition of *Streptococcus mutans* strains in non-acidic conditions is intriguing. As, logically, expression of the MGC would be required for conferring a metabolic and potentially disease-suppressive phenotype, metatranscriptomics analysis of samples where putative polyacrolein accumulation is observed could be an interesting follow-up analysis in the future to test the hypothesis of the involvement of this MGC in its production. Additionally, more detailed chemical analysis of the putative polyacroleins is required to confirm their structural identity. Nonetheless, this analysis illustrates how BiG-MAP analysis, especially when combined with complimentary omics data such as metabolomics, can generate concrete and relevant hypotheses about microbiome-associated phenotypes that can be tested in the laboratory.

**Figure 3.**
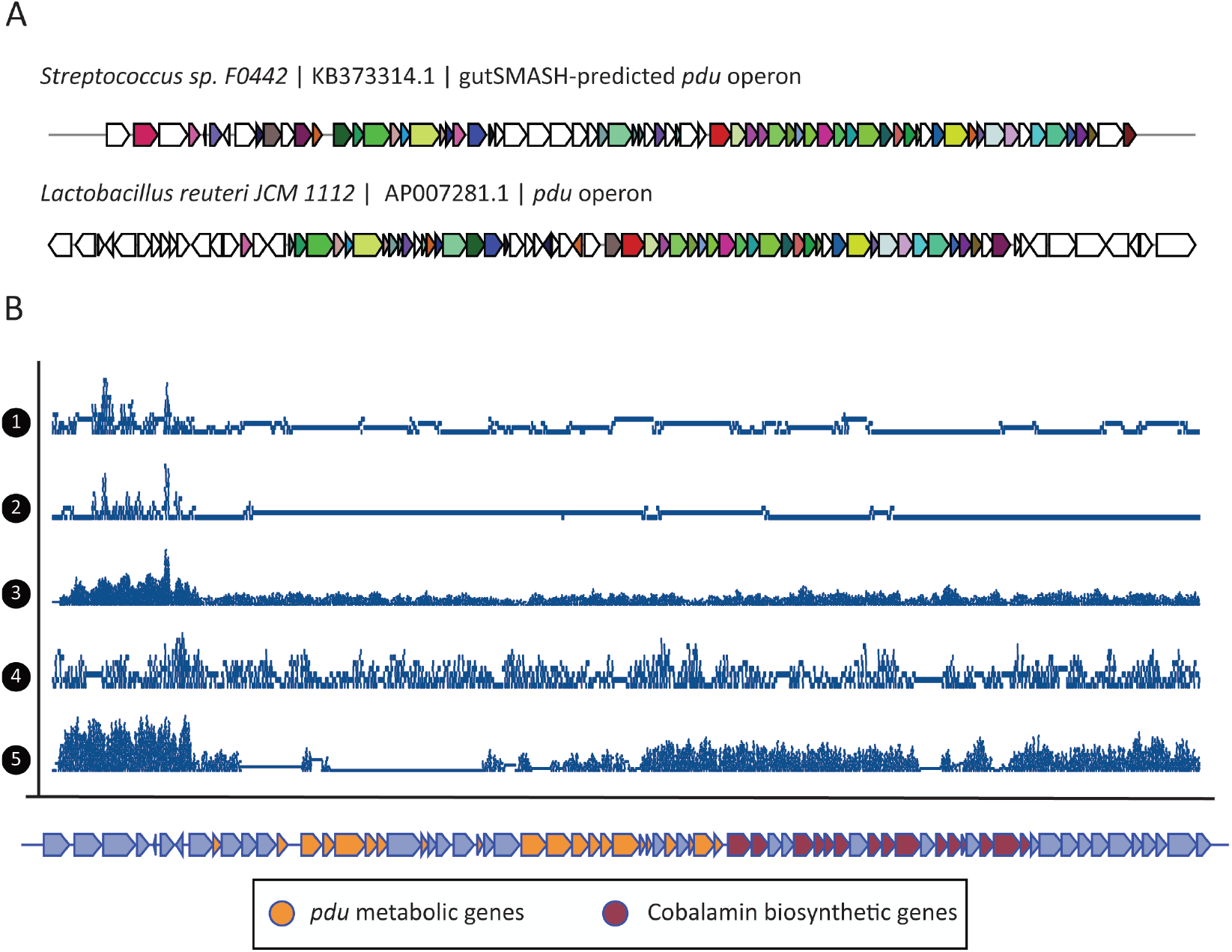
Detection of a *pdu* / cobalamin operon in healthy oral metagenomes. (A) MultiGeneBlast comparison between the *pdu* operon found in *Streptococcus sp. F0442* by gutSMASH and the characterized one from *Lactobacillus reuteri* (AP007281). (B) Read coverage of five randomly chosen healthy metagenomes along the gutSMASH-predicted *pdu* gene cluster. The coverage graphs, which were plotted using the Sushi R package (version 3.5.1)^44^, show that some samples (3 and 4) contain reads that cover the whole gene cluster, while in other samples, reads hardly cover the cluster (1 and 2) or only part of it (5).

Another example of a gene cluster that has been found relevant in the oral cavity is the *muc* operon, which has been shown to be responsible for the production of tetramic acid, which is known to inhibit the colonization of commensal bacteria in the oral cavity. This gene cluster encodes a hybrid between a polyketide synthase and nonribosomal peptide synthetase (PKS/NRPS)^32^. In order to further test this association and assess the abundance of the *muc* operon in the oral cavity, a collection of 170 *Streptococcus mutans* genomes collected from Tang *et al*^32^ and Liu *et al*^45^ was run through antiSMASH^10^, which predicted a total of 1,849 BGCs. After obtaining 41 representative gene clusters with BiG-MAP.family module, reads from the 47 oral microbiome metagenomes were mapped onto the predicted gene clusters and further processed using BiG-MAP.map and BiG-MAP.analyse subsequently. From the results, two gene clusters were found to be significantly differentially abundant between healthy and disease samples when using the fitZIG model: an NRPS from *Streptococcus mutans* N29 and the *muc* operon from *Streptococcus mutans* 14D. The *muc* operon from this strain shows high similarity to the one characterized by Tang *et al*.^32^ (Cumulative Blast bit score of 9,056) (see Fig. 4A). However, the mean read core coverage in both groups is low; 0.283 in healthy and 0.372 in caries-associated samples, which imply the presence of some of the *muc* operon genes but not the complete gene cluster (see Fig. 4B). Nonetheless, within both groups we see that some samples have reads mapping to the complete gene cluster, with coverages values close to 1. When filtering out samples with coverage values < 0.5; leaving only 6 samples in each group, the mean coverage rises to 0.803 in healthy and 0.991 in disease. This is because there are nine healthy samples that have a core coverage value of 0 and also five disease samples that do not have reads mapping to the core genes of the *muc* operon. Interestingly, depending at which stage you check which group is more enriched with this gene cluster—either before or after normalization and depending on which differential abundance test you apply—one group or the other seems to have higher counts. The average abundance of raw RPKM counts in healthy is 16854.08 compared to 12815.69 in disease samples. After being normalized, healthy samples have on average 11.22 RPKM counts/sample, slightly lower than the disease group that has 11.28 RPKM counts/sample. When using the two available differential abundance testing methods, we see that when applying the fitZIG model the difference in abundance between healthy and disease samples is significant (more abundant in disease) but not when testing it with Kruskal-Wallis. This is illustrated in the fitZIG BiG-MAP output heatmap (Suppl. Fig. S3), which shows that despite the *muc* operon is significantly more abundant in disease samples, the abundance of this gene cluster across all samples is generally very similar. Therefore, despite finding this operon being more abundant in caries-prone samples when applying the fitZIG model, suggesting that indeed the *muc* operon plays a role in the caries development, the oral microbiota from healthy donors seem to also harbor this PKS/NRPS. Hence, the microbiota from healthy samples may have a mechanism to counteract the inhibiting effect of tetramic acid, or there might be a difference in expression of the gene cluster between healthy and diseased subjects.

**Figure 4.**
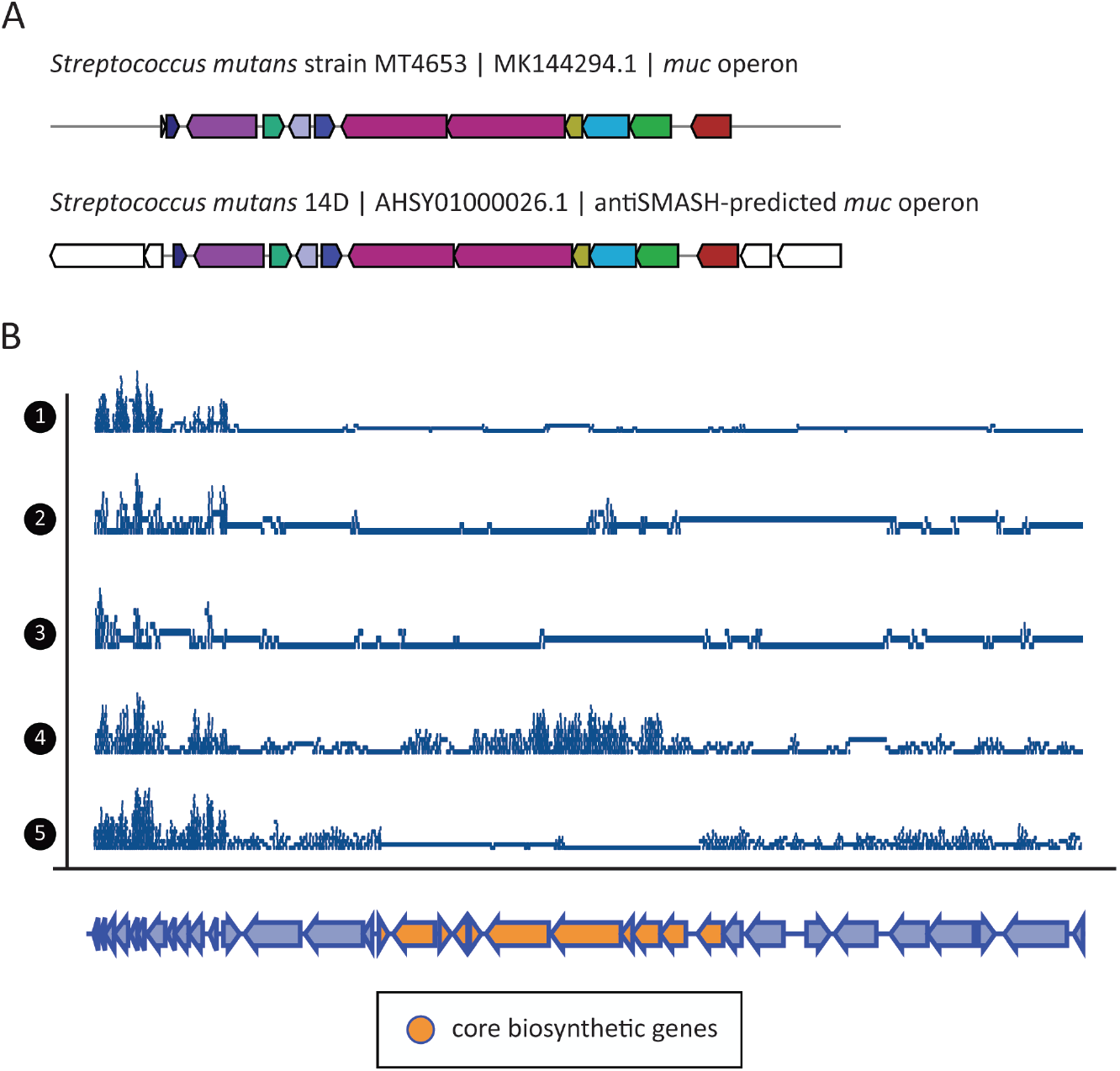
Detection of the *muc* operon in a subset of caries-associated samples. (A) MultiGeneBlast comparison between the *muc* operon characterized from *Streptoccocus mutans* strain MT4653.1 and the antiSMASH predicted one from *Streptococcus mutans* 14D. (B) Read coverage of five random chosen caries-related metagenomes along the antiSMASH predicted *muc* gene cluster. The coverage graphs, which were plotted using the Sushi R package (version 3.5.1)^44^, show that despite the fact that the *muc* operon is generally not very highly covered by reads from the randomly picked examples, some seem to truly contain for this operon, such as sample 4, where the core biosynthetic genes look to be abundant at sufficient levels. Full data pertaining all samples can be found in Fig. S3.

In addition, we also assessed the presence of the *muc* operon in the extended dataset that includes 96 metagenomic samples in total (see *Evaluating the presence of the muc operon in caries-associated metagenomes* Methods section). However, neither the *muc* nor the other BGCs predicted from the *Streptococcus* genomes were significantly more abundant in either group. This could be explained because within the 96 samples there are not only healthy or caries-associated metagenomes but also metagenomes from patients suffering from periodontitis and samples from a study that observes how a biofilm evolves over time; therefore, it might be that the community structure of all these samples differ quite a lot in terms of BGC content but also regarding the presence of *Streptococcus mutans*. All in all, our results suggest that the abundance of the *muc* operon is not very predictive for a healthy or disease state of the microbiome by itself, and other factors likely play (more) important roles.

## Conclusions

Overall, combining different omics datasets is a very useful approach to understand which microbes are doing what and poses a promising avenue to better understand complex biological processes. Here, we presented BiG-MAP, a command-line tool that it is able to profile the abundance and expression of a collection of gene clusters across metagenomic and metatranscriptomic data. Each of the steps in the BiG-MAP pipeline is robust, as demonstrated using simulated metagenomes. Indeed, BiG-MAP can discover interesting and relevant potential associations between genomic regions and phenotypes, which can guide experimental efforts to test MGC function. It is worth noting the usefulness of the gene cluster mapping coverage values, since they allow the user to discern between the real presence of predicted gene clusters of interest and spurious read mapping. Also, the associations that can be found using BiG-MAP strongly depend on the WGS data sequencing depth and sample size, as for instance in the examples described in our study, we found both gene clusters (*pdu-like* operon and *muc*) only significant in either dataset (reduced or extended one). Moreover, from the BiG-MAP output folders, which include raw and processed results, it is possible to extract valuable information, such as the differences within groups, distribution of reads across a gene cluster, raw and normalized RPKM counts, etc. Overall, we believe BiG-MAP will help researchers solving biologically complex questions by integrative multi-omics approaches, to obtain deeper insights into the relationships between microbial metabolic capacities and microbiome-associated phenotypes.

## Methods

### Code availability

BIG-MAP is implemented in Python 3 as a command line package. It consists of four modules: BiG-MAP.download, BiG-MAP.family, BiG-MAP.map, and BiG-MAP.analyse. The code is available at: https://github.com/medema-group/BiG-MAP together with documentation on how to install BiG-MAP and its dependencies and a short tutorial on how to run it.

### BiG-MAP.download: Data collection

This module allows to retrieve sequencing data present in the SRA database using the SRA toolkit (https://github.com/ncbi/sra-tools). To initially develop, test and validate this, we used an IBD cohort that contains metagenomic and metatranscriptomic data from 78 individuals, 21 suffering from UC, 46 individuals with CD, and 11 healthy samples^46^. These samples were retrieved using the SRA accession IDs under BioProject PRJNA389280 tool (see Suppl. Fig S1 and S2 generated from this dataset).

### BiG-MAP.family: Creating a non-redundant MGC representative collection

The family module uses as input a directory that contains the gene cluster prediction outputted by the antiSMASH^47^ or gutSMASH algorithms (https://github.com/victoriapascal/gutsmash). The predicted gene clusters are then subjected to a redundancy filtering step based on their mutual sequence similarity. For that, the protein sequences of the gene clusters are extracted and used as input for MASH^20^ sketch, which creates sketches from the raw sequences. The sketches are then used to calculate the distances between sequences using MASH dist. The resulting tab-delimited file with the pairwise distance comparisons is used to group together gene clusters with above a 0.8 default similarity cut-off (see Figure 2). Next, to pick the best representative of each group, medoids are computed (see formula below). For this, a distance matrix is created comparing all distances between pairs of gene clusters; the one with minimal cumulative distance value is picked as representative of that group. Additionally, the selected gene clusters are subjected to another round of clustering using BiG-SCAPE^18^, to group gene clusters into GCFs at a 0.3 similarity cut-off (default value), from which a random representative is picked.

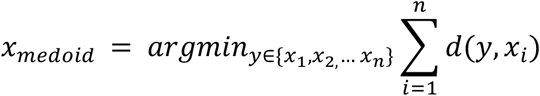

If metatranscriptomes will be used in the BiG-MAP.map module, an additional step is performed to set an expression baseline. For this, the protein sequences of the genomes whose gene clusters form the non-redundant representative gene cluster collection are scanned using hmmsearch (hmmsearch version 3.1b2) for five housekeeping-coding proteins: DNA gyrase A (PF00521), DNA gyrase B (PF00204), Recombinase A (PF00154), DNA directed RNA polymerase A (PF01000), and DNA directed RNA polymerase B (PF00562). The selection of these Pfam domains was based on the findings by Rocha *et al*.^48^ that these housekeeping genes show highly stable expression across samples. Next, the gathered protein sequences are also used as queries in the mapping module to align metatranscriptomic reads to gene clusters.

### BiG-MAP.map: mapping reads to a non-redundant gene cluster collection

This module relies on Bowtie2^21^ (version 2.3.4.3) to align reads to a given sequence. From the reference gene cluster sequences selected by the medoid calculation, Bowtie index files are created. Next, Bowtie2 aligns reads to these index files that by default uses the fast alignment mode. The resulting alignment is stored in SAM format and converted to BAM format to later be parsed by SAMtools^49^ (version 1.9). The alignments are then sorted by leftmost coordinates, the aligned reads are counted and corrected by GCF and gene cluster size consecutively. Later, the corrected raw counts are converted to TPM counts (Transcripts Per Kilobase Million) and consecutively to RPKM (Reads Per Kilobase Million) counts to account for sequencing depth.

Another functionality that was added in this module was to compute the read coverage of each gene cluster using the coordinates in the sorted BAM files. To do so, the sorted alignment files are converted to bedgraphs using BEDtools^50^ (v2.28.0), that allow to estimate the number of covered bases of each cluster (*coverage*) by subtracting the number of non-covered bases (*ncb*) to the length of each cluster (*cl*) as indicated in the formula below.

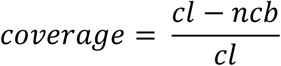

The same procedure is followed to compute the RPKM counts and the coverage of the core genes within a gene cluster, which strictly considers the core metabolic genes within each gene cluster. This information is taken from the antiSMASH/gutSMASH (or any other “SMASH” related algorithm) Genbank output files that flag the key coding genes that are needed for the synthesis of a given molecule. Once the core genes are identified, the alignment information concerning them is retrieved using SAMtools. Next, in the same manner as RPKM are computed for the whole gene clusters, reads aligned to the core region are pulled out, counted and corrected to finally get the RPKM counts. To perform the coverage calculation, the locations of the core genes are extracted from the bedgraph to evaluate the coverage score using the aforementioned formula.

### BiG-MAP.analyse: Normalization of RPKM counts and finding differentially expressed/abundant MGCs

In order to account for sparse high-throughput sequencing RPKM are normalized using Cumulative Sum Scaling (CSS) from the R Bioconductor package MetagenomeSeq^22^. BiG-MAP offers two different statistics to account for differentially abundant/expressed gene clusters, the parametric zero inflated gaussian distribution mixture model (ZIG-models) that assumes normal distribution of values or the non-parametric Kruskal-Wallis test. Relatively small changes in gene cluster abundance/expression are expected thus, ZIG-model values are adjusted with log2 fold-change that ultimately helps fitting the model to a log-normal distribution. Alternatively, Kruskal-Wallis can be run on the normalized RPKM counts, which allows to assess whether the distribution of ranks for one group significantly differs from the distribution of ranks for the other group. Additionally, FDR correction is applied to correct for multiple hypothesis testing. Finally, heatmaps are produced to visualize the results using the Seaborn python package (https://github.com/mwaskom/seaborn).

### Testing BiG-MAP performance using a mock community

To test BiG-MAP performance, 101 bacterial genomes were randomly chosen from the CGR collection^24^. Thus, the gutSMASH-predicted MGCs from each genome were used as ground truth (https://github.com/victoriapascal/gutsmash, version 0.8, github commit stamp: 569e860). Next, paired-end reads were generated with a mean read length of 100 bp from the 101 CGR bacterial genomes using Grinder v0.5.3^51^. Two different read coverage thresholds were used (0.5x and 0.05x) in combination with four different community structures: uniform, linear, power-law and exponential. Both the MGCs and the simulated reads were used as input for BiG-MAP, which was run ranging the MASH similarity thresholds between 10-100% in intervals of 10% along the eight different Bowtie2 alignment modes. From each individual run, true positive, false positive and false negatives rates were calculated to evaluate the precision and recall, which was ultimately used to compute the harmonic mean of precision and recall, also known as the F1-score. The results were plotted in a heatmap using the ComplexHeatmap package in R^52^.

### Assessing the *pdu* operon abundance by surveying different oral metagenomic samples

To find possible leads on metabolic perturbances between healthy and caries-related samples, the processed mass spectra (MGF format) and metabolomics feature tables from Aleti, G. *et al.*^30^ were downloaded from GNPS-MassIVE^34^ accession ID MSV000081832 to perform re-analysis. Feature-based Molecular Networks^53^ were run using GNPS release version 21 (https://gnps.ucsd.edu/ProteoSAFe/status.jsp?task=ef4f64542ab24a7fb0802ceacbcfa071, https://gnps.ucsd.edu/ProteoSAFe/status.jsp?task=9c95754d1fdc42b4a43b16919c398ecd). The resulting molecular family information together with the metabolite feature tables and sample information (metadata) were loaded into PALS (https://pals.glasgowcompbio.org/app/)^33^, to identify metabolite families differing in activity between healthy and caries-related samples. From the results, three out of seven candidate metabolites in one differentially expressed molecular family showing clear different abundance patterns between healthy and caries samples were further examined using GNPS MASST (https://masst.ucsd.edu)^36^, the ChemCalc MF finder^54^, and PubChem^55^, leading to the putative annotation of polyacrolein-related metabolites in healthy samples, which may be produced from a *pdu-like* operon that requires the presence of the cobalamin biosynthetic genes (see Supplementary material for further information).

For the analysis of the *pdu* operon and its presence in the oral microbiome, 1,440 oral bacteria genomes were downloaded from the HOMD collection (http://www.homd.org/?name=GenomeList&link=GenomeList&type=all_oral). Next, these genomes were used as input for gutSMASH (version 0.8). The comparison between the two *pdu* operons from *Lactobacillus reuteri* (AP007281) and *Streptococcus sp. F0442* (GCA_000314795.2) was done using MultiGeneBlast^56^. Next, all predicted gene clusters were used as input for the BiG-MAP family module. At the same time, the oral metagenomics datasets were downloaded using the BiG-MAP.download module by providing the SRA accession IDs associated to the PRJNA478018, PRJNA396840, and PRJNA398963 BioProject IDs. Once the metagenomes were downloaded, BiG-MAP.map was run using the output of the family module and the metagenomic reads in fastq format. Finally, the RPKM counts were normalized, processed and visualized using BiG-MAP.analyse.

### Evaluating the presence of the *muc* operon in caries-associated metagenomes

AntiSMASH was used to predict BGCs from a total of 170 *Streptococcus mutans* genomes reported in Tang *et al*^32^ and Liu *et al*^45^. Within the predicted BGCs, the *muc* operon was found and compared to the *muc* operon characterized by Hao *et al*.^57^ using MultiGeneBlast^56^. The predicted BGCs were then used as input for the BiG-MAP.family module. Both, the representative BGCs and metagenomic reads were then used as input in the subsequent BiG-MAP.map mapping module using the metagenomes from the following three BioProjects: PRJNA478018, PRJNA396840, and PRJNA398963. Finally, the raw mapping counts were normalized and further processed and visualized using BiG-MAP.analyse.

### Data availability

The supporting information for this article can be found in the Supplementary material and in the Zenodo repository (https://zenodo.org/) with the following DOI: 10.5281/zenodo.4320501. The metabolomics data used for reanalysis is available from GNPS-MassIVE accession ID MSV000081832.

## Acknowledgements

We thank Daria Zuzanna Świgoń, Arno Hagenbeek, Sarah van den Broek, Jeanine Boot and Robert Koetsier for preliminary results on the *pdu* operon, which provided us the lead to further explore these datasets. We also acknowledge the guidance provided by Rens Holmer in the early stage of this study and Dr Madeleine Ernst for her help in locating the relevant files of the relevant metabolomics data files from the Aleti *et al*. study.

## Funding information

This work was supported by the Chan-Zuckerberg Biohub (M.A.F.), and the U.S. Defense Advanced Research Projects Agency’s Living Foundries program award HR0011-15-C-0084 (M.A.F. and V.P.A.) and an ASDI eScience grant (ASDI.2017.030) from the Netherlands eScience Center (J.J.J.v.d.H. and M.H.M.).

## Conflicts of interest

MHM is a co-founder of Design Pharmaceuticals and a member of the scientific advisory board of Hexagon Bio. M.A.F. is a co-founder and director of Federation Bio, a co-founder of Revolution Medicines, and a member of the scientific advisory board of NGM Biopharmaceuticals.

## Notes

### Competing Interest Statement

The authors have declared no competing interest.

