## Supplementary material for "BiG-MAP: an automated pipeline to profile metabolic gene cluster abundance and expression in microbiomes"

### Supplementary Methods

#### Metabolomics analyses

GNPS Molecular Networking was done as follows:

A molecular network was created with the Feature-Based Molecular Networking (FBMN) workflow on GNPS (https://gnps.ucsd.edu). The data was filtered by removing all MS/MS fragment ions within +/- 17 Da of the precursor m/z. MS/MS spectra were window filtered by choosing only the top 6 fragment ions in the +/- 50 Da window throughout the spectrum. The precursor ion mass tolerance was set to 0.02 Da and the MS/MS fragment ion tolerance to 0.02 Da. A molecular network was then created where edges were filtered to have a cosine score above 0.65 and more than 5 matched peaks. Further, edges between two nodes were kept in the network if and only if each of the nodes appeared in each others respective top 10 most similar nodes. Finally, the maximum size of a molecular family was set to 100, and the lowest scoring edges were removed from molecular families until the molecular family size was below this threshold. The analogue search mode was used by searching against MS/MS spectra with a maximum difference of 200.0 in the precursor ion value. The library spectra were filtered in the same manner as the input data. All matches kept between network spectra and library spectra were required to have a score above 0.7 and at least 6 matched peaks.

GNPS Molecular Networking job of mass spectrometry data: https://gnps.ucsd.edu/ProteoSAFe/status.jsp?task=9c95754d1fdc42b4a43b16919c398ecd

MASST searches were performed for a number of features that showed typical C3H4O mass differences using the default settings. A single spectrum search was completed using the online workflow (https://ccms-ucsd.github.io/GNPSDocumentation/) on the GNPS website (http://gnps.ucsd.edu). The data was filtered by removing all MS/MS fragment ions within +/- 17 Da of the precursor m/z. MS/MS spectra were window filtered by choosing only the top 6 fragment ions in the +/- 50 Da window throughout the spectrum. The precursor ion mass tolerance was set to 2.0 Da and a MS/MS fragment ion tolerance of 0.5 Da. The library spectra were filtered in the same manner as the input data. All matches kept between input spectra and library spectra were required to have a score above 0.7 and at least 6 matched peaks.

Feature ID 3779, precursor m/z 680.4799:

https://gnps.ucsd.edu/ProteoSAFe/result.jsp?task=71831f592f27496faf16c62bc16b1b69&view=view_all_datasets_matched

Feature ID 153, precursor m/z 722.4900:

https://gnps.ucsd.edu/ProteoSAFe/result.jsp?task=79bc074e6e274f05ac4d3a273c50681a&view=view_all_datasets_matched

Feature ID 126, precursor m/z 663.4528 :

https://gnps.ucsd.edu/ProteoSAFe/result.jsp?task=8742b10dda2a4e29bc405fdc34aa1aa6&view=view_all_datasets_matched

**Supplementary Figure S1**

Illustration of BiG-MAP analysis module output when using metagenomics data. In this case, for testing and developing purposes, the CGR genomes were used as input for gutSMASH. The resulting predictions were used as input together with the metagenomics samples from Schirmer *et al.* (PRJNA389280). The significant differential abundance of 11 gene clusters across Crohn disease (CD) samples and healthy (non-IBD) using Kruskal Wallis are shown in the heatmap. The more abundant a gene cluster is, the more yellow it is shown in the heatmap. Next to the heatmap, the bar chart represents the abundance log2 fold-change values. On the far most right, the dots represent the coverage values computed by BiG-MAP to show how evenly the reads map along the whole gene cluster, in blue for the CD samples and in orange for the healthy.

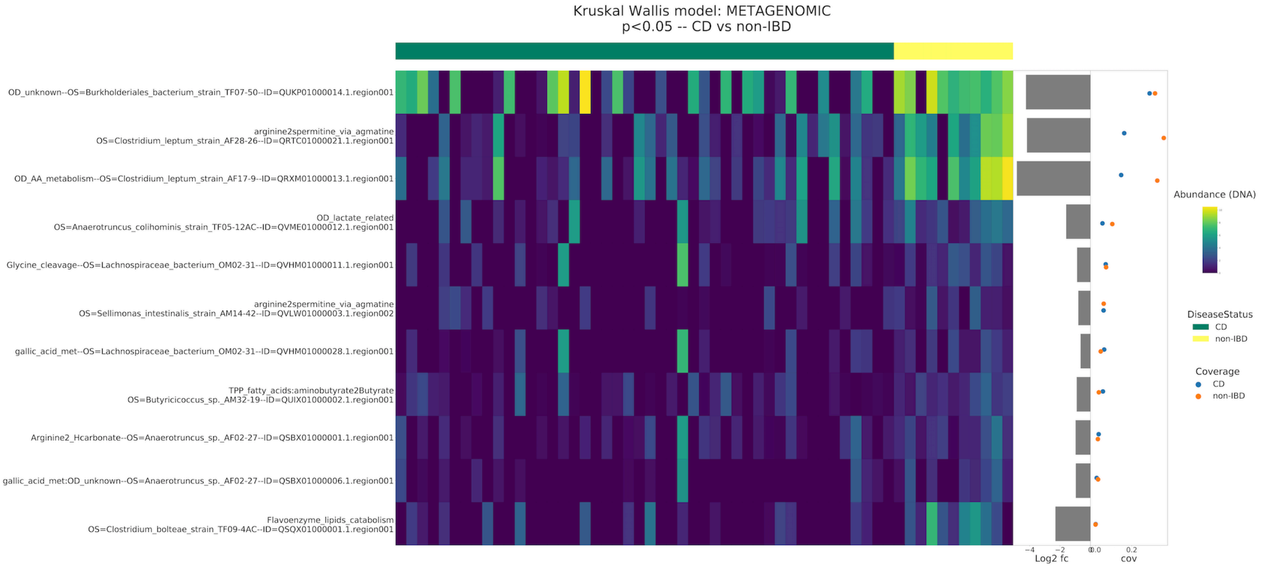


**Supplementary Figure S2**

Illustration of BiG-MAP analysis module output when using metatranscriptomic data. In this case, for testing and developing purposes, the CGR genomes were used as input for gutSMASH. The resulting predictions were used as input together with the metatranscriptomes from Schirmer *et al*. The significant differential expression of 8 gene clusters across Crohn disease (CD) samples and healthy (non-IBD) using Kruskal Wallis are displayed in the heatmap. The higher the expression of a gene cluster is, the more yellow it is shown in the heatmap. Similarly to Figure S1, the log2 fold-change and coverage values are also depicted. When analyzing metatranscriptomic data, it is also included a heatmap with the expression values of five housekeeping genes that allows to set an expression baseline.

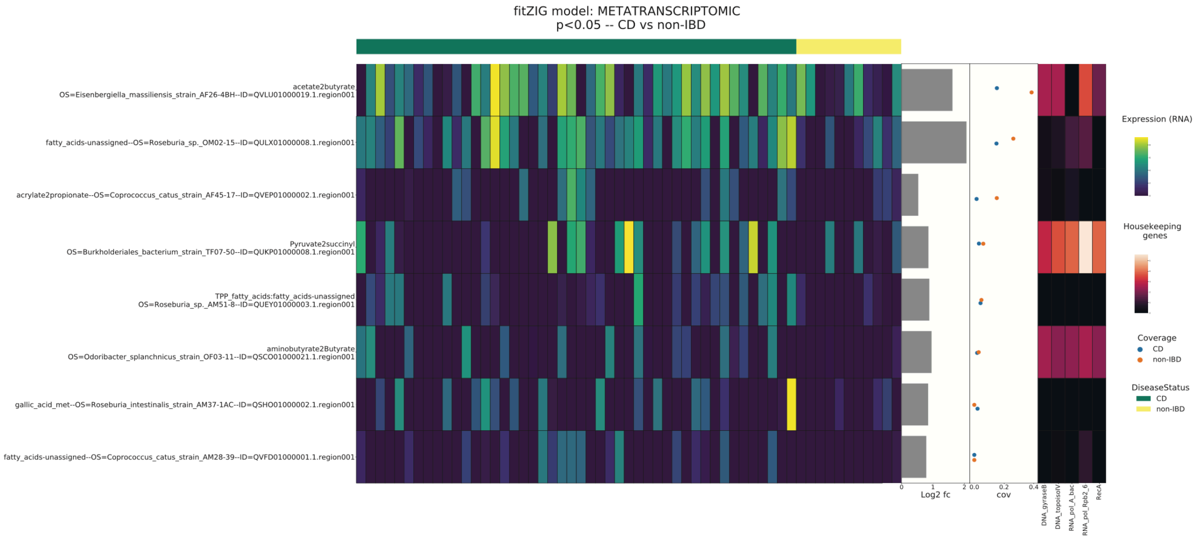


**Supplementary Figure S3**

The *muc* operon (AHSY01000026.1.region001), predicted by antiSMASH, is significantly enriched in disease samples when applying the FitZIG model. Despite the significance, the difference in abundance between both groups is minimal as the similar coloring depicts and the log2 fold-change bar chart evinces. The plot shows another gene cluster, also an NRP, found significantly enriched in disease samples The heatmap has been produced by BiG-MAP.analyse module using the 47 oral microbiome metagenomes.

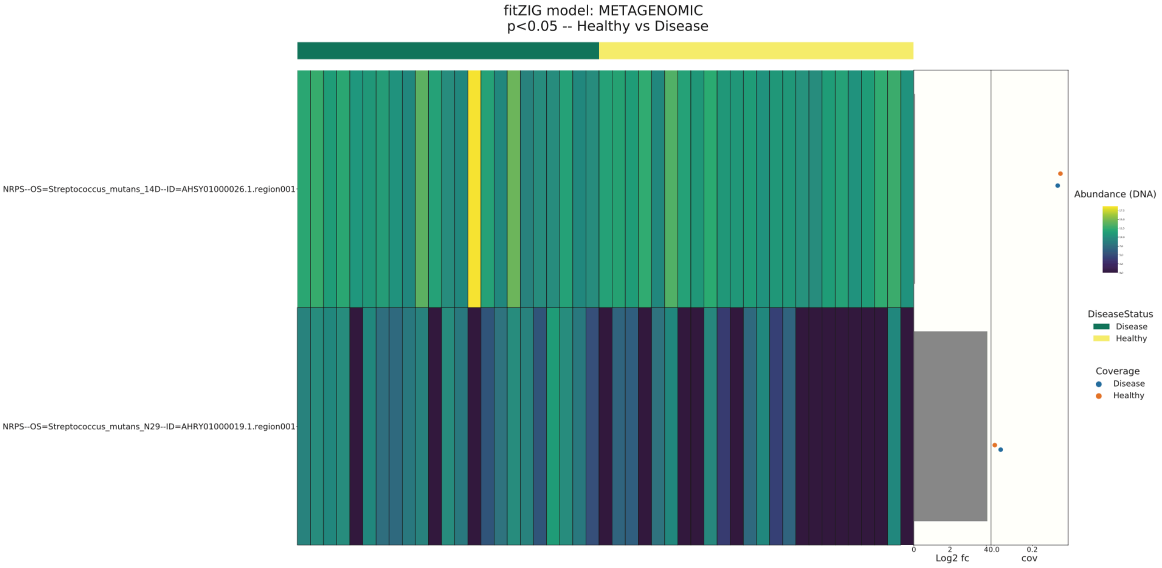
fitZIG model: METAGENOMIC
p<0.05 -- Healthy vs Disease
NRPS--OS=Streptococcus_mutans_14D--ID=AHSY01000026.1.region001
NRPS--OS=Streptococcus_mutans_N29--ID=AHRY01000019.1.region001
0
2
40.0
0.2
Log2 fc
cov
Abundance (DNA)
DiseaseStatus
Disease
Healthy
Coverage
Disease
Healthy
